# Screening and Development of Constitutively Synergistic Combination Drug Formulations for T Cell Acute Lymphoblastic Leukemia

**DOI:** 10.1101/2022.09.21.508196

**Authors:** James M Kelvin, Dan Y Zhang, Evelyn K Williams, Samuel G Moore, Lacey A Birnbaum, Henry Zecca, Xiaodong Wang, Juhi Jain, Min Qui, Nathan T Jui, Haian Fu, Yuhong Du, Melissa L Kemp, Wilbur A Lam, Deborah DeRyckere, Douglas K Graham, Erik C Dreaden

**Author notes:** equal contribution (James M Kelvin, Dan Y Zhang, Evelyn K Williams).

## Abstract

Advances in multiagent chemotherapy have led to recent improvements in overall survival for patients with acute lymphoblastic leukemia (ALL); however, a significant fraction do not respond to frontline chemotherapy or later relapse with recurrent disease, after which long-term survival rates remain low. To address the challenge of developing new, effective treatment options for these patients, we conducted a series of high-throughput combination drug screens to identify chemotherapies that synergize in a lineage-specific manner with MRX-2843, a small molecule dual MERTK and FLT3 kinase inhibitor currently in clinical testing for treatment of relapsed/refractory leukemias and solid tumors. Using experimental and computational approaches, we found that MRX-2843 synergized strongly – *and in a ratio-dependent manner* – with vincristine chemotherapy to inhibit T-ALL cell expansion and, based on these findings, we developed multiagent lipid nanoparticle formulations of these drugs that not only constitutively maintained ratiometric drug synergy following T-ALL cell delivery, but also improved anti-leukemic activity following drug encapsulation. To determine the clinical relevance of these combination drug formulations and the therapeutic impact of ratiometric drug synergy, we compared the efficacy of lipid nanoparticles comprising synergistic, additive, and antagonistic ratios of MRX-2843 and vincristine, and observed that trends in *in vitro* synergy were directly recapitulated in primary T-ALL patient samples. Together, these findings present a systematic approach to high-throughput combination drug screening and multiagent drug delivery that maximizes the therapeutic potential of combined MRX-2843 and vincristine in T-ALL. This broadly generalizable approach could lead to the development of constitutively synergistic combination products for the treatment of cancer and other diseases.

## INTRODUCTION

Combination approaches to cancer therapy have greatly improved treatment outcomes since their inception in the mid-1950s [1-3] and today, a majority of cancer patients receive some form of multiagent therapy [4] that can encompass a wide range of drug types including small molecules, proteins, nucleic acids, viral vectors, or engineered cells. In addition to diversity of size and structure, these agents also vary widely in their route or frequency of administration and, as a result, large time-dependent fluctuations in both tissue and plasma drug concentration – and therefore drug ratio – are commonly observed following the administration of combination therapies [5-7].

Drug synergy – the observation of combined drug effects that exceed the sum of component drug effects – is a common basis for the use of multiagent therapies in cancer, infectious diseases, and neurological disorders. Yet, despite the strong therapeutic potential of synergistic multiagent approaches, prior large-scale screening studies suggest that the identification of synergistic drug pairs is in fact rare (4-10%) [8], while others indicate that synergistic effects – when they do occur – are highly ratio-dependent, whereby some ratios of particular drugs may be supra-additive (*i*.*e*. synergistic) while others are additive or antagonistic (*i*.*e*. sub-additive) [9-11].

Recent approaches to maximize the therapeutic potential of multiagent cocktails have aimed to co-deliver therapeutics in a manner that constitutively maintains drug synergy [12]; these include nanometer-scale drug carriers that deliver agents in a time-staggered [13, 14] or ratiometric [15-19] fashion, as well as those that harness synthetic lethal gene interactions [20] or target adaptive drug resistance mechanisms *a priori* [21, 22]. Liposomal cytarabine and daunorubicin (CPX-351, Vyxeos^®^) is one such example and was originally discovered through *in vitro* screens that identified synergistic cell growth inhibition at a 5:1 mole ratio of cytarabine:daunorubicin [23, 24]. By constitutively maintaining ratiometric drug synergy between these two agents via lipid nanoparticle formulation, CPX-351 improved treatment outcomes relative to free combination chemotherapy in both mouse models and clinical trials and is now FDA-approved for the treatment of both adult and pediatric patients with leukemia [25-27].

The receptor tyrosine kinase, MERTK, is ectopically expressed in a majority of acute myeloid leukemias (AML) and approximately 50% of T cell acute lymphoblastic leukemias (T-ALL) and we previously described key roles for the protein in leukemia cell survival and leukemogenesis [28-31]. More recently, we developed small molecule inhibitors that target MERTK and FLT3, a clinically-validated therapeutic target in AML [32]. These agents improve survival in murine ALL and AML models [33-35] and the lead compound, MRX-2843, is currently being tested in Phase 1/2 clinical trials (NCT03510104, NCT04762199, NCT04872478, NCT04946890). We hypothesized that the therapeutic potential of combined MERTK/FLT3 inhibition and chemotherapy for leukemia could be maximized via ratiometric drug screening and formulation. To this end, here we describe a novel and systematic approach to high-throughput combination drug screening and nanoscale drug delivery that maximizes drug synergy between MRX-2843 and cytotoxic chemotherapy. This novel, broadly generalizable approach to formulation discovery could lead to the development of constitutively synergistic combination products for the treatment of cancer and other diseases.

## RESULTS

Guided by prior reports indicating the potential for synergy between MERTK inhibition and vincristine or methotrexate [28, 31, 33] we developed a high-throughput drug screen in which we sought to identify ratiometric synergy between these compounds following cell exposure to drug interaction matrices comprising concentration gradients of MRX-2843, vincristine, and methotrexate **(Fig 1a)**. As an *in vitro* model, we selected a genotypically and phenotypically diverse set of leukemia cell lines spanning B- and pro-B ALL, as well as T- and early T cell precursor (ETP-) ALL, exposing each of these cell lines to drug interaction matrices comprising >530 discrete ratiometric drug combinations, and their equivalent single agents, in 384-well plates using high-throughput liquid handling robotics **(Fig 1b)**. Cell density was measured in quadruplicate via luminescent cell viability assay using Z’≥0.5 as a cutoff for data quality.

**Figure 1.**
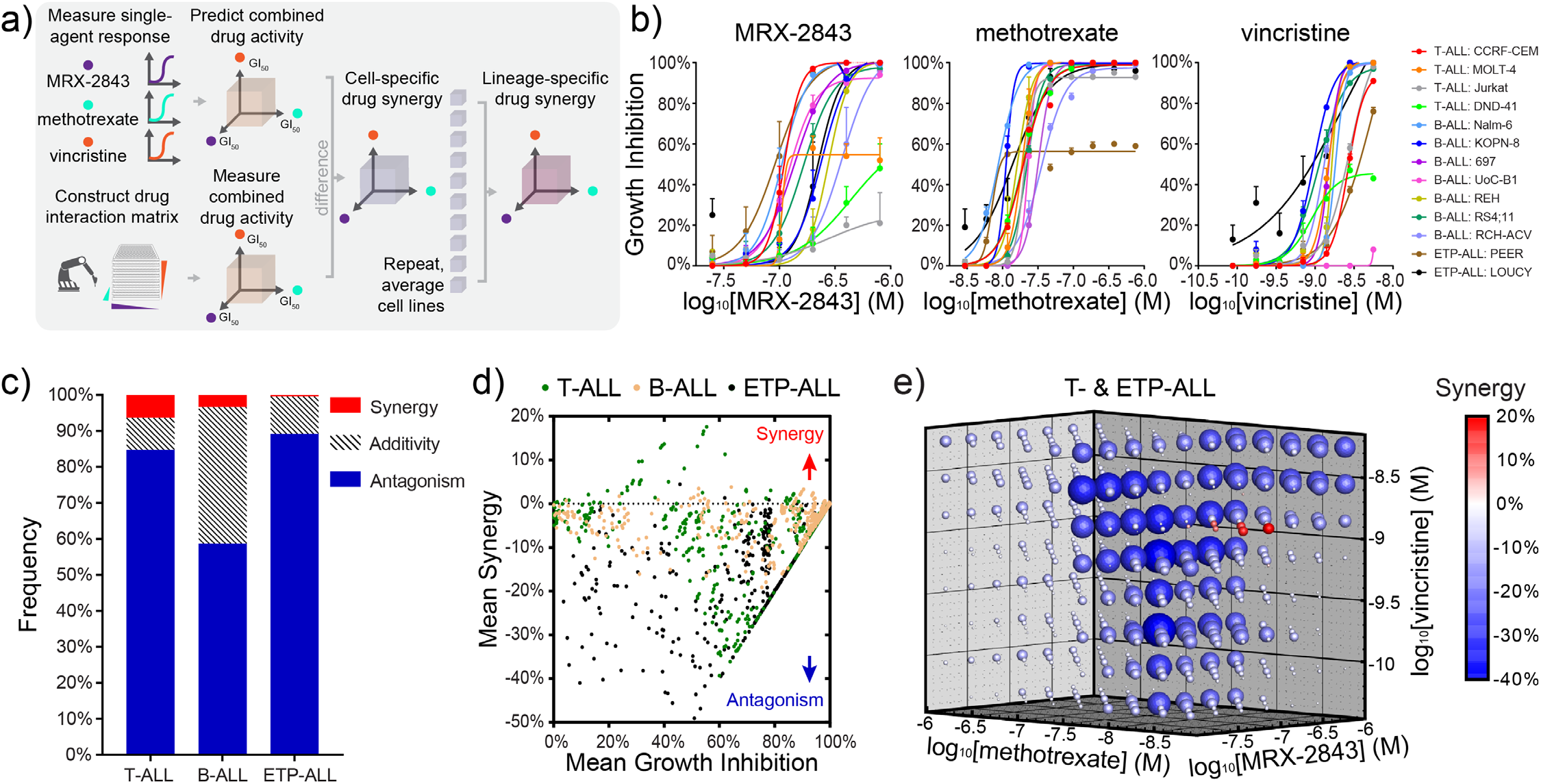
Combined MRX-2843 and vincristine chemotherapy synergize to inhibit T-ALL cell expansion in high-throughput combination drug screens. **a)** Illustration of a ratiometric drug screen combining the dual MERTK/FLT3 inhibitor, MRX-2843, with methotrexate and/or vincristine chemotherapy in a panel of (13) B- and T-ALL cell lines as measured by luminescent viability assay (72 h, Z’≥0.5). **b)** Single agent dose-response curves and **(c)** lineage-specific frequencies of combined drug synergy, additivity, or antagonism for 537 unique pairwise or triplet drug combinations as measured in parallel and assessed via the Response Additivity model. **d)** Scatter plots of mean drug synergy as a function of mean reduction in cell denstiy grouped by cell lineage. **e)** Drug synergy and antagonism is conserved across T- and ETP-ALL cell lines at distinct molar ratios of MRX-2843 and vincristine. Data in (b) represent mean ± SD of n=4 replicates. Synergy in (c-e) represent percent reduction in cell density above or below that predicted by the response additivity model with synergy (>1%) and antagonism (<1%).

Consistent with other large scale screening studies [9, 36, 37], we observed that synergy among these ratiometric combinations was rare (<3.4% overall) as assessed using the Response Additivity model of drug synergy **(Fig 1c,d)** [38]. Interestingly, drug synergy was largely absent when averaged over B lineage ALL cells **(Fig S1)**; however, we observed strong synergy *and* antagonism that was ratio-dependent across T lineage ALL cell lines **(Fig 1e)**. The consistency of these trends in synergy was confirmed using the Bliss Independence and Zero Interaction Potency synergy models [38] and the screening results were reproduced upon validation using an alternative 96-well assay format and new drug and cell stocks **(Fig S2a-c)**.

To guide the selection of a drug combination with maximum potency in T-ALL, we next determined whether synergy observed in our prior screen was principally attributable to either higher order (3-drug) or lower order (2- and 1-drug) effects. By comparing concordance between measured cell viabilities and those predicted from synergy models [39], we found that synergy from the prior drug screen was predominantly attributable to lower order, rather than higher order 3-drug interactions **(Fig 2a, S3)**. To confirm which among the three pairwise drug combinations was most synergistic, we conducted a microwell assay in which hydrogel-embedded Jurkat T-ALL cells were exposed to continuous gradients of each drug pair and assessed for cell viability. Comparing viability data to corresponding additive isoboles, we found that the combination of MRX-2843 and vincristine was most consistently synergistic **(Fig 2b, S2d**,**e)**.

**Figure 2.**
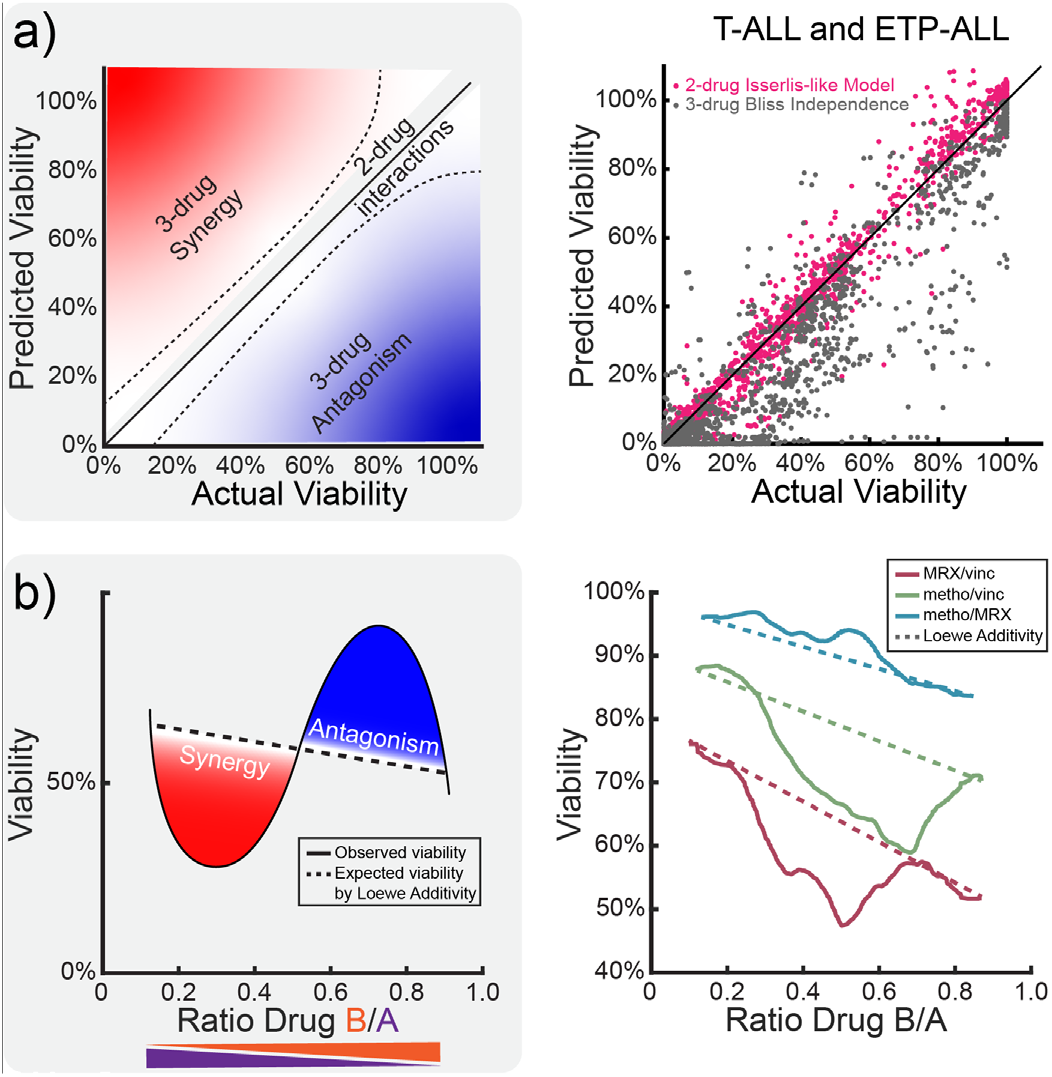
*In silico* and *in vitro* models prioritize pairwise drug synergy between MRX-2843 and vincristine among screening combinations. **a)** Comparison of higher order (3-drug) and lower order (2- and 1-drug) synergy models to high-throughput combination screening data indicates that experimentally observed drug synergy is predominantly attributable to pairwise drug interactions. **b)** Comparison of expected (dashed) and observed (solid) Jurkat T-ALL cell viability following exposure to continuous pairwise drug gradients demonstrating consistent synergy between MRX-2843 and vincristine. Data in (b) represent mean cell density (72 h) as measured by nuclear dye exclusion.

Having determined that MRX-2843 and vincristine act in a ratio-dependent manner to decrease T-ALL cell density, we next sought to develop drug formulations that both deliver and constitutively maintain ratiometric synergy between these compounds. Using a chemoinformatics-guided approach, we devised a pH-gradient based method of drug loading **(Fig 3a,b)** in which the compounds were simultaneously co-encapsulated within lipid nanoparticles composed of DSPC (1,2-distearoyl-sn-glycero-3-phosphocoline), DSPG (1,2-distearoyl-sn-glycero-3-phosphoglycerol, sodium salt), and cholesterol lipids. Using this approach, we synthesized drug formulations of MRX-2843 and vincristine that recapitulated drug ratios which we had previously identified as synergistic (17.9 - 71.4 mol:mol MRX:vinc), additive (4.5 - 8.9 mol:mol), or antagonistic (142.9 - 9,143.0 mol:mol) in their inhibition of T-ALL cell expansion (*referred to hereafter as Syn, Add, and Ant nanoparticles, respectively*). Characterization of the subsequent lipid nanoparticle formulations by dynamic light scattering, transmission electron microscopy, and liquid chromatography-mass spectrometry (LC-MS) indicated high uniformity (PDI 0.11 - 0.14) and efficient drug loading (5 - 6 wt%) using these methods **(Fig 3c,d, S4a-c)**.

**Figure 3.**
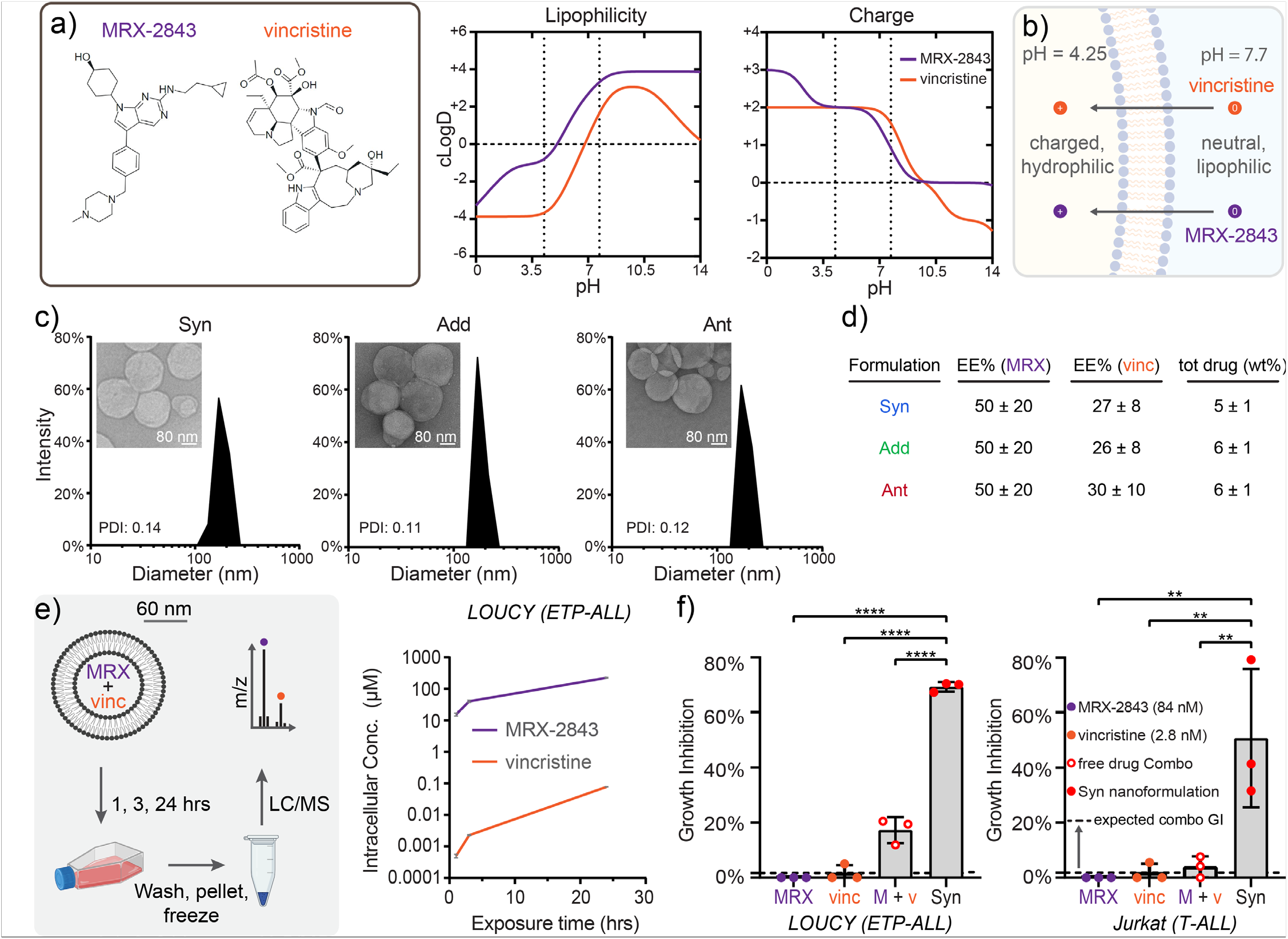
Lipid nanoparticles co-encapsulating MRX-2843 and vincristine constitutively maintain synergistic drug ratios following co-formulation and intracellular delivery. **a)** Structure and calculated pH-dependent lipophilicity or charge of MRX-2843 and vincristine. **b)** Illustration of a pH-gradient based method of luminal drug co-loading within liposomes. **c)** T-ALL tailored combination drug formulations encapsulating synergistic (Syn), additive (Add), and antagonistic (Ant) drug ratios illustrating nanometer-scale size and uniformity as measured by dynamic light scattering and (*inset*) transmission electron microscopy. **d)** Table of drug loading parameters (encapsulation efficiency (EE%) and loading capacity (tot drug (wt%)) as measured by LC-MS. **e)** Intracellular drug delivery kinetics over 24 h following treatment of Loucy ETP-ALL cells with Syn nanoparticles as measured by LC-MS. **f)** Comparison of cell densities following treatment (72 h) with Syn nanoparticles, as well as individual or combined free MRX-2843 and free vincristine in Loucy ETP-ALL or Jurkat T-ALL cells as measured by luminescent viability assay. Data in (f) represent mean ± SD of n=3 replicates from a single experiment, and the dashed line represents the expected additive value for combined free drugs assessed via the Response Additivity model. Differences were measured by one-way ANOVA, where * < 0.05, ** < 0.01, *** < 0.001, and **** < 0.0001.

To confirm that combination drug formulations of MRX-2843 and vincristine indeed achieved ratiometric drug delivery, we exposed Loucy cells to drug-loaded nanoparticles for varying times over 24 hours, then measured intracellular drug content via LC-MS (**Fig 3e**). We found that Syn nanoparticles not only achieved highly efficient delivery of MRX-2843 (their principal drug component) to T-lineage ALL cells, but that they also constitutively maintained intracellular MRX-2843 and vincristine ratios over time. To further characterize the therapeutic performance of Syn nanoparticles, we examined their impact on Loucy and Jurkat cell density via luminescent viability assay. *Strikingly*, drug synergy was even further enhanced in both cell lines following nanoparticle encapsulation, whereby growth inhibition far exceeded that of the already synergistic combination of free MRX-2843 and vincristine (Loucy, P<0.0001; Jurkat, P=0.0090) **(Fig 3f)**.

After demonstrating strong concordance between ratiometric synergy from free- and lipid nanoparticle-encapsulated MRX-2843 and vincristine, we investigated the clinical relevance of these combination drug formulations using primary pediatric T-ALL patient samples collected at time of diagnosis. Following optimization of culture conditions, mononuclear cells from bone marrow or blood were treated with liposomal MRX-2843, free vincristine, vehicle lipid nanoparticles, or Syn or Add nanoparticles that reflect optimal synergy or additivity in Loucy ETP-ALL cells **(Fig 4a)**. GI_50_ values of cell density, measured via luminescent viability assay, demonstrated greater potency for Syn and Add formulations in cell lines than in primary patient samples; however, the magnitude of growth inhibition observed in these primary samples was notably high (>58-78% for Syn, 45-91% for Add) **(Fig 4b,c)**, and both Syn and Add formulations recapitulated their expected synergism or additivity in these patient samples as compared to expectations based on the effects of liposomal MRX-2843 and free drug vincristine measured in parallel **(Fig 4d,e)**.

**Figure 4.**
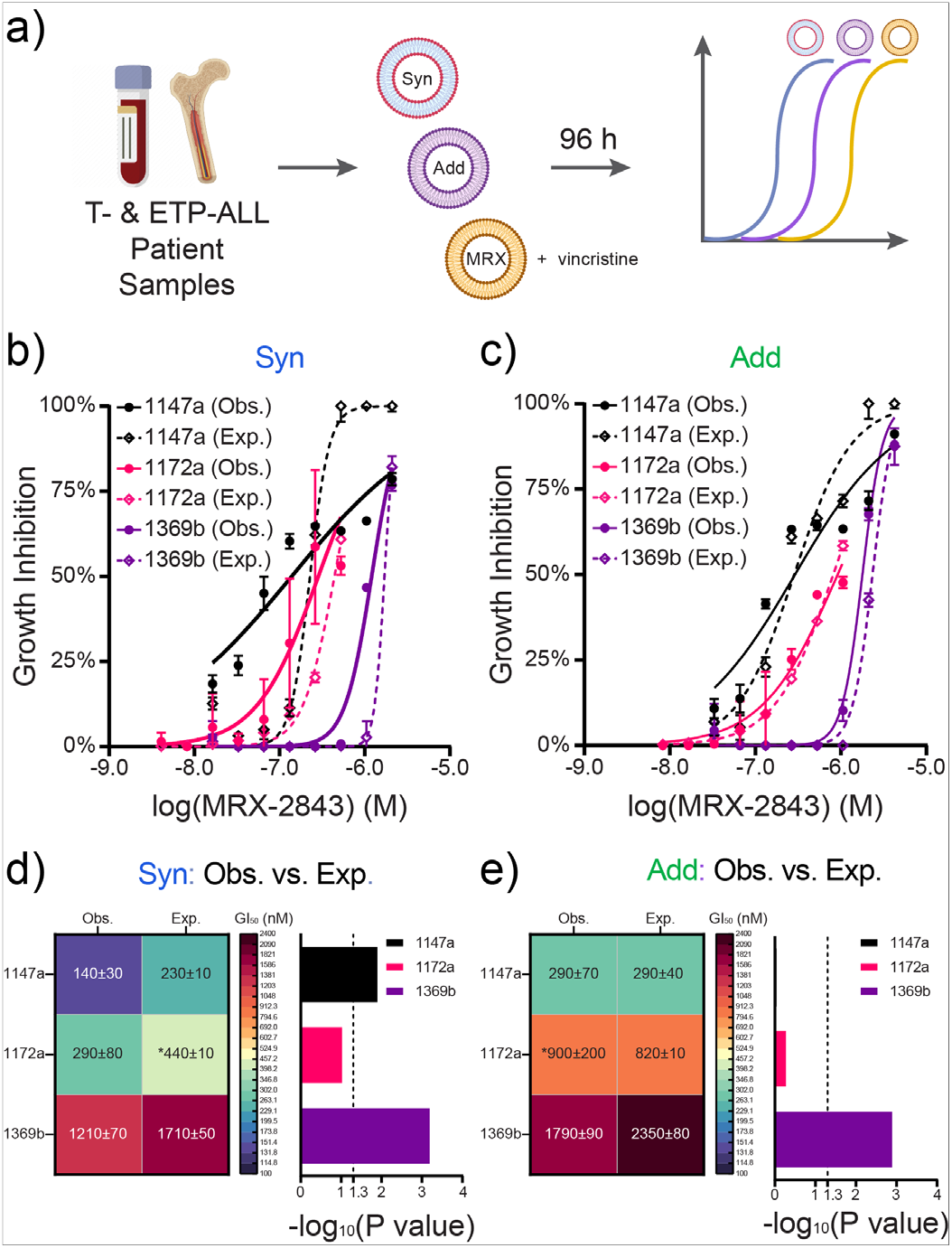
Comparative efficacy of ratiometric drug formulations in patient T-ALL and ETP-ALL samples. **a)** Approach to testing primary T-ALL and ETP-ALL samples against ratiometric nanoformulations. **b)** Comparative potencies of observed (Obs.) and expected (Exp.) Syn and **c)** Add nanoparticles in T-ALL or ETP-ALL samples obtained from patients at initial diagnoses as measured by luminescent viability assay (96 h). Dose response curves represent observed (Obs.) or expected (Exp.) response additivity based on the sum of liposomal MRX and vincristine free drug monotherapies. **d)** Syn nanoparticles achieve significant reductions in GI50 compared to expectation, while **e)** Add nanoparticles overlap with expectation. Data in (b,c) represent mean and standard deviation of GI values scaled against the concomitant MRX-2843 doses in the nanoparticles. Expected dose response curves were calculated with the Response Additivity model of drug synergy. Heat maps in (d,e) show GI_50_ ± standard error (nM) and associated log transformed P values of unpaired parametric t tests. Asterisks (*) in (d,e) denote extrapolated GI_50_ values. All Obs. dose responses were run in triplicate, while Exp. dose responses were run either in duplicate (1172a, bone marrow derived near ETP-ALL) or triplicate (1147a, bone marrow derived T-ALL; 1369b, blood derived T-ALL) in a single experiment.

## DISCUSSION

Here we address the urgent and unmet need to develop new, increasingly potent therapies for patients with relapsed/refractory T-ALL. While advancements in chemotherapy dose-intensification and patient risk-stratification have led to gradual and significant improvements in patient survival for T-ALL in recent years, a substantial proportion of patients have treatment refractory disease or later relapse. As a result, mortality rates for pediatric and adult T-ALL patients are 20% and >50%, respectively [40]. Patients among the ETP-ALL subtype experience even higher rates of induction therapy failure that both necessitate intensified treatments associated with increased toxicity and increase their likelihood for bone marrow transplant [41-45]. To address the challenge of improving outcomes following salvage therapy in these patients, we devised a systematic approach to maximize drug synergy between small molecule anticancer drug combinations and to tailor these effects to a specific disease indication, in this case T-ALL.

Using a novel, high-throughput combination drug screen, we found that combined treatment with the tyrosine kinase inhibitor, MRX-2843, and the cytotoxic chemotherapy, vincristine, synergized in a ratio-dependent manner to inhibit T-ALL cell expansion, a finding that is significant because (i) vincristine is a mainstay of frontline induction and consolidation therapies for T-ALL [46] and (ii) both MRX-2843 targets are frequently actionable in T-ALL. For example, in our previous studies, approximately 50% of pediatric T-ALL patient samples ectopically expressed MERTK [28] and 50% of both pediatric and adult samples were sensitive to treatment with a MERTK/FLT3 inhibitor [47]. Similarly, approximately 15% and 35% of pediatric and adult ETP-ALLs, respectively, carry activating *FLT3* mutations that confer sensitivity to FLT3 inhibition [48, 49].

Given the dual targets of MRX-2843 and the multifarious effects of MERTK on oncogenic signaling pathways [50], the precise mechanism(s) by which MRX-2843 synergizes with vincristine in T-ALL are speculative; however, pharmacologic inhibitors of several pathways and proteins that are known to be downstream targets of MERTK signaling also synergized with vincristine, suggesting their potential as mediators of synergy. MERTK promotes expression of the anti-apoptotic protein BCL-xL in ALL and other cancer cells and can inhibit expression of the related protein, BCL-2, in some circumstances [31, 51]. Gossypol, a BCL-2 and BCL-xL inhibitor, synergized with vincristine in non-Hodgkin’s lymphoma cells [52]. Likewise, inhibition of BCR/ABL whose downstream signaling, like MERTK, converges on the MAPK pathway, synergized with vincristine against ALL cells [53]. Lastly, inhibitors of PI3K – positively regulated by MERTK signaling – synergized with vincristine in T-ALL cell culture and murine models [54]. While mechanistic studies to further characterize the drug synergy observed here are underway, together these studies suggest that the *combined* role of MERTK in regulating apoptosis, as well as cell growth/proliferation and survival signaling, may contribute to its role in promoting resistance to vincristine.

Based on the findings from our screen and subsequent analyses indicating ratiometric drug synergy between MRX-2843 and vincristine, we developed multiagent lipid nanoparticles closely related in lipid composition to FDA-approved products such as AmBisome and Vyxeos, both of which are comprised of phosphatidylcholine, phosphatidylglycerol, and cholesterol [55]. To further enhance the feasibility of clinical-scale manufacturing, we also devised a method to co-load defined ratios of MRX-2843 and vincristine using a gradient-based method which is notable in its typically high encapsulation efficiency, scalability, and reproducibility, as well as its utilization to manufacture FDA-approved products, Onivyde and Doxil [56, 57]. Given the safe and effective clinical use of these lipid excipients, together with a commercially viable method of manufacturing, we anticipate high translational potential of liposomal MRX-2843 and vincristine in the future.

Having identified synergy between MRX-2843 and vincristine and developed lipid nanoparticle formulations co-loading these compounds, we further examined the extent to which combination formulations maintain ratiometric drug synergy following cell delivery. Indeed, the Syn combination formulation achieved fixed *intracellular* drug ratios of MRX-2843 and vincristine *in vitro*. These findings support the success and importance of the pH gradient-based loading method described here. Drugs contained within the lumen of lipid vesicles are typically released less rapidly than those that reside in the lipid bilayer [14], where lipophilic compounds often reside following passive drug loading or incomplete gradient-based loading [56, 57]. The comparable intracellular drug release profiles between MRX-2843 (clogD_7.4_=3.07) and vincristine (clogD_7.4_=1.14) observed here thus suggest that our loading method was not only successful in luminal drug entrapment, but that it was also critical to maintaining ratiometric drug synergy following cell delivery. Further, the intracellular drug concentrations achieved were well in-excess of their corresponding GI_50, 72h_ values in T-ALL cells. Together, these findings support both the therapeutic utility and clinical relevance of our described approach to co-formulating MRX-2843 and vincristine.

In addition to maintaining synergistic drug ratios following cell delivery, the synergistic anti-leukemic activity observed in cell cultures treated with the free drug combination was further enhanced following nanoparticle co-encapsulation, a phenomenon that we and others have previously observed with other drug pairs and carriers [15]. While studies to elucidate the mechanisms by which drug co-encapsulation *further* improves synergy between MRX-2843 and vincristine are ongoing, such effects may be attributable in part to nanoparticle-accelerated intracellular drug transport, augmented target engagement whereby the intracellular kinase domains of both MERTK and FLT3 (enzymatic IC_50_∼ 1.3 and 0.64 nM [34]) are exposed to drug concentrations in excess of those achieved via passive drug diffusion, or differential plasma/serum protein binding. Further, in addition to prominent cell-surface expression, MERTK has also been reported to co-localize with early endosomes [58] where lipid nanoparticles frequently accumulate following cellular uptake [59]; thus, nanoparticle-mediated drug delivery may augment the magnitude, duration, and/or location of protein inhibition in order to further enhance drug synergism.

Lastly, we confirmed the clinical relevance of these novel, T-ALL-tailored combination drug formulations in primary blood and bone marrow samples collected from patients with T-ALL at initial diagnosis. While primary T-ALL samples were more refractory to treatment with MRX-2843 and vincristine than established cell lines, as reported for other MERTK/FLT3 inhibitors [60], both drug synergy and additivity were maintained by Syn and Add drug formulations, respectively, following *ex vivo* treatment. These findings are significant in that the delivery of drug carriers and transfection reagents to primary T lymphocytes is notoriously difficult [61]. Given the synergistic potency and efficient formulation and delivery described here, this work may serve to guide future clinical studies of these or related drug formulations incorporating MRX-2843 and/or other anticancer agents for the treatment of ALL.

## MATERIALS AND METHODS

### Cell Lines and Culture

Nalm-6, KOPN-8 and UoC-B1 were obtained from the lab of Dr. Christopher Porter (Emory University). 697, REH, RCH-ACV, RS4;11, CCRF-CEM, DND-41, Jurkat, MOLT-4, LOUCY and PEER were obtained from Dr. Deborah DeRyckere and Dr. Douglas Graham (Emory University). Cells were cultured in RPMI 1640 with L-Glutamine supplemented with 10-20% heat-inactivated fetal bovine serum or as recommended by the original supplier (DSMZ, ATCC). All cells were cultured at 37 °C in a 5% CO_2_ humidified atmosphere and tested regularly for mycoplasma.

### High-Throughput Combination Drug Screening

MRX-2843 (obtained under MTA from Meryx Inc.) and methotrexate (Sigma-Aldrich) were reconstituted in dry DMSO and stored in checkerboard array within 384-well polypropylene plates (VWR) covered in sterile adhesive plate film prior to use. Vincristine sulfate (Cayman Chemical) was solvated in molecular biology grade water (VWR) on the day of experimentation. 15,000 viable cells per well were dispensed into white polystyrene 384-well microplates (Greiner Bio-One) using a Multidrop Combi Reagent Dispenser (Thermo Fisher). Vincristine was subsequently dispensed onto these cell-containing microplates. MRX-2843 and methotrexate were then transferred to these cell + vincristine microplates using a Beckman NX Liquid Handler (Beckman Coulter) at the Emory Chemical Biology Discovery Center. Cell- and drug-containing plates were then centrifuged for 5 minutes at 135 x g (Eppendorf 5810R), then incubated for 72 hours at 37°C in 5% CO_2_ humidified atmosphere with gas permeable plate sealing film (VWR). All wells had a DMSO concentration of 0.5% v/v with drugs MRX-2843 (25 – 800 nM), methotrexate (2.9375 – 752 nM), and vincristine (0.0875 – 5.6 nM). After 72 hours, viable cell numbers were assessed using CellTiter-Glo 2.0 Assay (Promega) and the Integra Viaflo Assist Pipetting Platform (Integra) according to manufacturer’s instructions. Luminescence was measured using a SpectraMax iD3 plate reader (Molecular Devices) and viability data were background corrected to empty wells containing DMSO/water/media and normalized to positive control cells treated with DMSO. All experiments were performed in quadruplicate using the following metrics as criteria for acceptability: CV<20% (positive and negative controls), SNR≥5 (positive controls), Z’≥0.5 (per plate). Drug synergy was calculated as described previously using the Response Additivity model and reported as %GI beyond the expected additive value, nominally classified using >1% and <-1% synergy as cutoffs for synergistic and antagonistic effects, respectively. Alternative calculations of drug synergy were performed as described previously using the Bliss Independence [62] and Zero Interaction Potency (ZIP) [63] models.

### Microwell Assay

Jurkat cells were embedded in 3 mg/mL collagen hydrogels (Corning) and were exposed simultaneously to overlapping concentration gradients of free drug or solvent controls for 72 hours. Following drug exposure, cell viability was assessed using viability dyes Calcein AM (ThermoFisher) and Propidium Iodide (ThermoFisher), and nuclei were stained using Hoechst 33342. Images were acquired using a Keyence BZ-X800 Fluorescence Microscope. Cell segmentation was performed using CellProfiler (v3.1, Broad Institute, Cambridge, MA), and cells were classified as live or dead based on mean per cell intensity measurements done in Matlab (MATLAB 2021a, Mathworks, Natick, MA). A rolling ball averaging method was then used to determine average viability at each drug combination dose ratio. Expected viabilities were computed based on Loewe Additivity isobolograms of single drugs at 10% of the respective maximum doses whereby the line of additivity is assumed to be a linear combination between the maximum effect at the highest concentrations of drug 1 and drug 2. Deviation from the line of additivity towards lower viability indicates synergy.

### 2- and 3-drug Effect Models

To determine whether synergy emerged from 2-drug or 3-drug interactions, we calculated pairwise and 3-drug combination synergy and antagonism using both lower order and higher order predictive models. For the lower order model, an Isserlis-like formula developed by Wood et al. [39] was used,

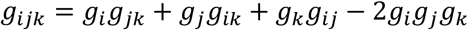

which when correlated with measured effects may indicate whether synergistic drug interactions can be explained by single and pairwise drug responses. For the higher order model, Bliss Independence was similarly used to assess correlations between measured and predicted values.

### Screening Validation

High-throughput screening (HTS) results were validated using an alternate assay plate format and new drug stocks and cell lots. Briefly, vincristine sulfate and MRX-2843 were dissolved in molecular biology grade water (VWR) on the day of experimentation, then serially diluted in cell media to obtain 0.01% v/v of water in all media dilutions. 45,000 viable Jurkat cells were then dispensed into drug/media-containing white polystyrene 96-well µClear microplates and handled as described in the above HTS experiments.

### Lipid Nanoparticle Formulation

DSPC (NOF), DSPG (NOF), and cholesterol (Sigma) were combined at a 7:2:1 mole ratio from lipid stocks solvated to 25 mg/mL in chloroform (DSPC, cholesterol) or 5 mg/mL chloroform:methanol (5:1 v/v, DSPG). Solvent from the lipid mixture was removed via rotary evaporation under vacuum at 30° C (Rotavapor R-100; Buchi) and the resulting lipid cake was further dried under house vacuum overnight. Lipid films were rehydrated via rapid addition of 250 mM ammonium sulfate buffer (pH 4.25) followed by high power cup horn sonication at 60° C (Q700; Qsonica). Multilamellar vesicles were then extruded ten times at (60° C) through 0.08 µm polycarbonate membranes (Nuclepore; Whatman) supported by polyester drain discs (230600; Whatman) using a high pressure N_2_ extruder (Liposofast LF-50; Avestin). Ammonium sulfate buffer was then exchanged for phosphate buffer (pH 7.7) via centrifugal diafiltration (100kDa Amicon) or tangential flow filtration (Krosflo KR2i; Spectrum) and unilamellar vesicles were then 0.45 µm sterile filtered (cellulose acetate, VWR).

To prepare drug-loaded lipid nanoparticles, MRX-2843 (10.0 mg/mL in molecular biology grade water) and vincristine (sulfate) (1.0 mg/mL in molecular biology grade water) were mixed at various molar ratios with freshly prepared, concentrated gradient liposomes. Drug and lipid nanoparticle mixtures were then agitated at 700 rpm (60° C) for 1 hour (Mixer HC; USA Scientific). Drug-loaded liposomes were dialyzed twice in phosphate buffered saline using 100kDa MWCO dialysis bags (Float-A-Lyzer G2; Spectrum Laboratories).

### UPLC-MS Analysis of Liposomal Drug Concentrations

Concentrations of MRX-2843, vincristine (sulfate), and DSPC were measured by LC-MS using a Vanquish Horizon UPLC (ThermoFisher Scientific) fitted with a Waters Corporation ACQUITY UPLC BEH C18 column (2.1×100 mm, 1.7 µm particle size) coupled to a high-resolution accurate mass Orbitrap ID-X Tribrid mass spectrometer. The chromatographic method for sample analysis involved elution with 20:80 MeCN:water with 10 mM ammonium formate and 0.1% formic acid (mobile phase A) and 10:90 MeCN:IPA and 0.1% formic acid with 10 mM ammonium formate (mobile phase B) using the following gradient program: 0 min 100% A; 0.5 min 100% A; 2 min 80% A; 8 min 0% A (curve 3); 10.9 min 0% A; 11 min 100% A; 12 min 100%A. The flow rate was set at 0.4 mL/min. The column temperature was set to 50 °C, and the injection volume was 2 µL.

Alternatively, the analytes were measured using a Waters Corporation Cortecs UPLC T3 column (2.1×150mm, 1.6 µm particle size). The chromatographic gradient for MRX-2843 and vincristine (sulfate) involved elution with water with 0.1% formic acid (mobile phase A) and MeCN with 0.1% formic acid (mobile phase B) using the following gradient program: 0 min 80% A; 2.8 min 50% A; 3.2 min 3% A; 5 min 3% A; 5.1 min 80% A; 5.5 min 80% A. The flow rate was set at 0.6 mL/min. The column temperature was set to 40 °C. The chromatographic method for DSPC involved elution with water with 0.1% formic acid (mobile phase A) and 90:10 IPA:MeCN with 0.1% formic acid and 10mM ammonium formate (mobile phase B) using the following gradient program: 0 min 40% A; 2.2 min 3% A; 5.4 min 3% A; 5.5 min 40% A; 7 min 40% A. The flow rate was set at 0.25 mL/min. The column temperature was set to 40 °C. The injection volume was 2 µL.

The Orbitrap ID-X is a tribrid spectrometer that utilizes quadrupole isolation with dual detectors, an orbitrap and an ion trap, with a maximum resolving power of 500,000 full width at half maximum (FWHM) at *m/z* 200 and mass accuracy of <1 ppm with use of an internal calibrant. The HESI source was operated at a vaporizer temperature of 275 ºC, a spray voltage of 3.5 kV in positive mode, and sheath, auxiliary, and sweep gas flows of 40, 8, and 1 (arbitrary units), respectively. Full MS was collected with 30,000 resolution at a range of 150-1500 mz.

UPLC-MS^2^ experiments were performed by acquiring mass spectra with targeted MS/MS (tMS^2^) acquisition. The MRX-2843 precursors, 489.3, 389.2, and 245.17 mz, were selected with a 0.8 mz isolation window, activated with 35% HCD, and the product ions were analyzed in the ion trap. The vincristine precursors, 825.4 and 413.2 mz, were selected with a 0.8 isolation window, activated with 35% CID, and the product ions were analyzed in the ion trap. The DSPC precursor, 790.6 mz, was selected with a 0.8 mz isolation window, activated with 30% HCD, and the product ions were analyzed in the orbitrap at 30,000 resolution. Data processing was performed with Thermo Scientific Xcalibur Version 4.3.73.11. The MSMS transitions were integrated, and the data was exported to excel. The transitions for each adduct were summed and quantified from standard curves.

### Lipid Nanoparticle Characterization

Hydrodynamic size and polydispersity were obtained via dynamic light scattering (DLS) (DynaPro, Plate Reader III; Wyatt Technologies) using 3 second acquisitions and averaged over 10 runs. Nanoparticle morphology was imaged via transmission electron microscopy (TEM) at 80 kV using a Hitachi HT-7700 instrument. Liposomes were diluted in ultrapure water and applied to charge coated formvar/carbon coated copper grids (400 mesh, Electron Microscopy Sciences) for 15 minutes followed by washing for 2 seconds with ultrapure water. Samples were then negatively stained with phosphotungstic acid for 20 seconds and washed again for 2 seconds with ultrapure water.

### Intracellular Drug Measurements

LOUCY cells were exposed to Syn nanoparticles under normal culture conditions, then cells were washed 3x in PBS and counted prior to snap-freezing on dry ice. Frozen cell pellets were prepared for LC-MS by mixing in 500 µL IPA and 50 µL of 500 µm glass beads. Samples were then homogenized in a Tissuelyzer II at 30 Hz for 5 minutes and the supernatant was transferred to separate microcentrifuge vials. The original samples in glass beads were washed twice with 100 µL of 80% MeOH, with each wash additive to the IPA supernatant. The same pipette tip was used for all supernatant transfer. The cellular extractions were then dried by vacuum centrifugation and reconstituted in 250 µL of 80% MeOH. Reconstitution was assisted by sonication for 5 minutes. The samples were centrifuged at 21,100 rcf prior to being transferred to LC vials for LC-MS analysis as described above.

### In vitro Syn Nanoformulation Dose Responses

LOUCY and Jurkat T-ALL cells were treated with dose-matched free drug monotherapies, the combination of free drugs, or Syn nanoparticles. Free drug and nanoparticles were dissolved in PBS vehicle (0.05% v/v) in media on white 96-well polystyrene µClear Microplates. 45,000 viable cells (Countess IIl Invitrogen) were added per well and cultured for 72 hours. Cells were then treated with CellTiter-Glo 2.0 (Promega) and analyzed for luminescence with a SpectraMax iD3 (Molecular Devices). Data represent technical triplicate measurements from a single experiment. % GI and synergy scores were calculated as described above.

### Primary Sample Responses to Ratiometric Nanoformulations

Viably frozen, de-identified blood and bone marrow samples collected from patients with T-ALL at initial diagnosis were obtained from the Aflac Cancer and Blood Disorders Center Leukemia and Lymphoma Biorepository at Children’s Healthcare of Atlanta. Cells (12,500 per 384-well) were cultured in cytokine-sera, consisting of 20 ng/mL human recombinant IL-7, IGF-1, and SCF (StemCell Technologies) in 90% RPMI + 10% heat inactivated FBS + 0.1 mM beta-mercaptoethanol (Sigma) [64-66] and exposed in triplicate to Syn or Add nanoparticles for 96 hours. Expected GI values were interpolated from dose response curves (GraphPad Prism) of free vincristine and lipid nanoparticle MRX-2843 using the Response Additivity model of drug synergy. Each well contained 0.5% v/v PBS/media. Positive controls were cells treated with PBS/media, and negative controls contained PBS/media but no cells. Following incubation, cultures were treated with CellTiter-Glo 2.0 and relative viable cell numbers were measured as described above.

### Statistical Analyses and Software

Differences between lipid nanoparticle and free drug dose responses in cell lines were measured by one-way ANOVA with Tukey’s multiple comparisons test. Observed and expected dose response GI_50_s for primary samples were compared by unpaired parametric t test. All synergy computations for dose responses were modeled as nonlinear 4-parameter logistic functions using GraphPad Prism. Multidimensional synergy plots were produced in Tecplot 360. Heat maps for primary sample potencies were rendered in OriginPro 2022. Chemoinformatics analysis and experimental diagrams were obtained using Chemicalize (Chemaxon) and BioRender software, respectively.

## SUPPORTING INFORMATION

B-ALL lineage-specific drug synergy, screening validation, cell line-specific low and higher-order drug synergy models, and drug loading and reproducibility.

## ACKNOWLEDGEMENTS

This work was supported in part by research grants from CURE Childhood Cancer (DD), No More Kids With Cancer (DKG, DD), the National Institutes of Health Research Training Program in Immunoengineering (T32EB021962), and the Coulter Department of Biomedical Engineering. We are also grateful for assistance from the Pediatric General Equipment Core, the Robert P. Apkarian Integrated Electron Microscopy Core, Georgia Institute of Technology’s Systems Mass Spectrometry Core Facility, and the Emory Chemical Biology Discovery Center. Patient samples were provided by the Aflac Cancer and Blood Disorders Leukemia and Lymphoma Biorepository at Children’s Healthcare of Atlanta; other investigators may have received specimens from the same subjects. The content here is solely the responsibility of the authors and does not necessarily represent the official views of the organizations acknowledged herein.

## AUTHOR CONTRIBUTIONS

J.M.K., X.W., N.T.J., H.F., Y.D., M.L.K., W.A.L., D.D., D.K.G., and E.C.D. designed research; J.M.K, D.Y.Z., E.K.W., S.G.M., L.A.B., H.Z., J.J., and M.Q. performed research or analyzed data; J.M.K., Y.D., M.L.K., W.A.L., D.D., D.K.G., and E.C.D. wrote or edited the manuscript.

## COMPETING INTERESTS

D.K.G. is a founder and serves on the Board of Directors of Meryx, Inc. D.D. and D.K.G. are equity holders in Meryx, Inc. J.M.K., J.J., D.D., D.K.G., and E.C.D. are inventors on a patent related to this work describing combination drug screening, formulation, and treatment.

