## Supplementary Information for "Screening and Development of Constitutively Synergistic Combination Drug Formulations for T Cell Acute Lymphoblastic Leukemia"

### SUPPORTING INFORMATION

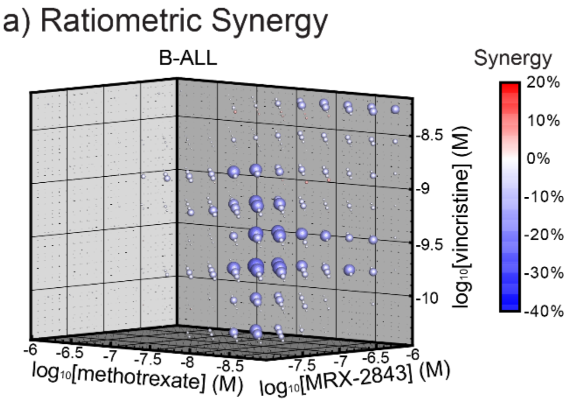


**SI Fig. 1. Neither synergy nor antagonism are conserved among B-ALL cells treated with varying concentrations of MRX-2843, methotrexate, and vincristine. a)** Mean ratiometric responses in B-ALL cell lines (n=7). Synergy represents percent reduction in cell density above or below that predicted by the Response Additivity model.

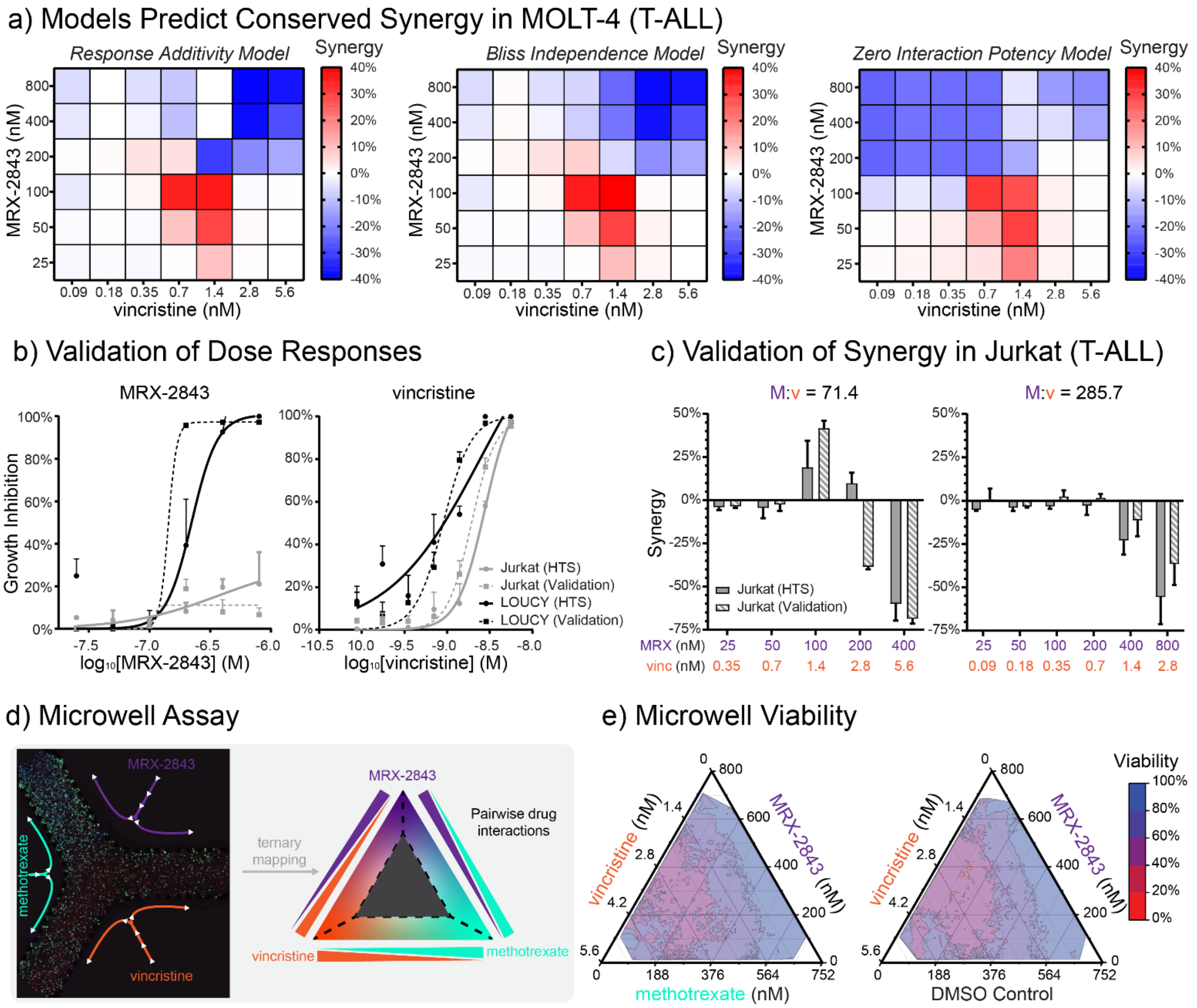


**SI Fig. 2. Alternative computational models and assay formats validate MRX-2843 and vincristine synergy. a)** Comparative analysis of ratiometric drug synergy in screening data from MOLT-4 T-ALL cells as calculated via Response Additivity, Bliss Independence, and Zero Interaction Potency models. Validation of **(b)** single-agent dose response and **(c)** combination drug synergy as measured via luminescent viability assay using an alternative (96-well) assay format and independent cell and drug stocks. **d)** Microwell assay of collagen-embedded Jurkat cells exposed to a continuous drug gradient of MRX-2843, methotrexate, and vincristine. **e)** Observed cell viability following exposure of Jurkat T-ALL cells to continuous drug gradients. Data in (b,c) represent mean ± SD of n=4 (HTS) or n=6 (Validation) replicates. Contours in (e) are averaged between 3 experiments.

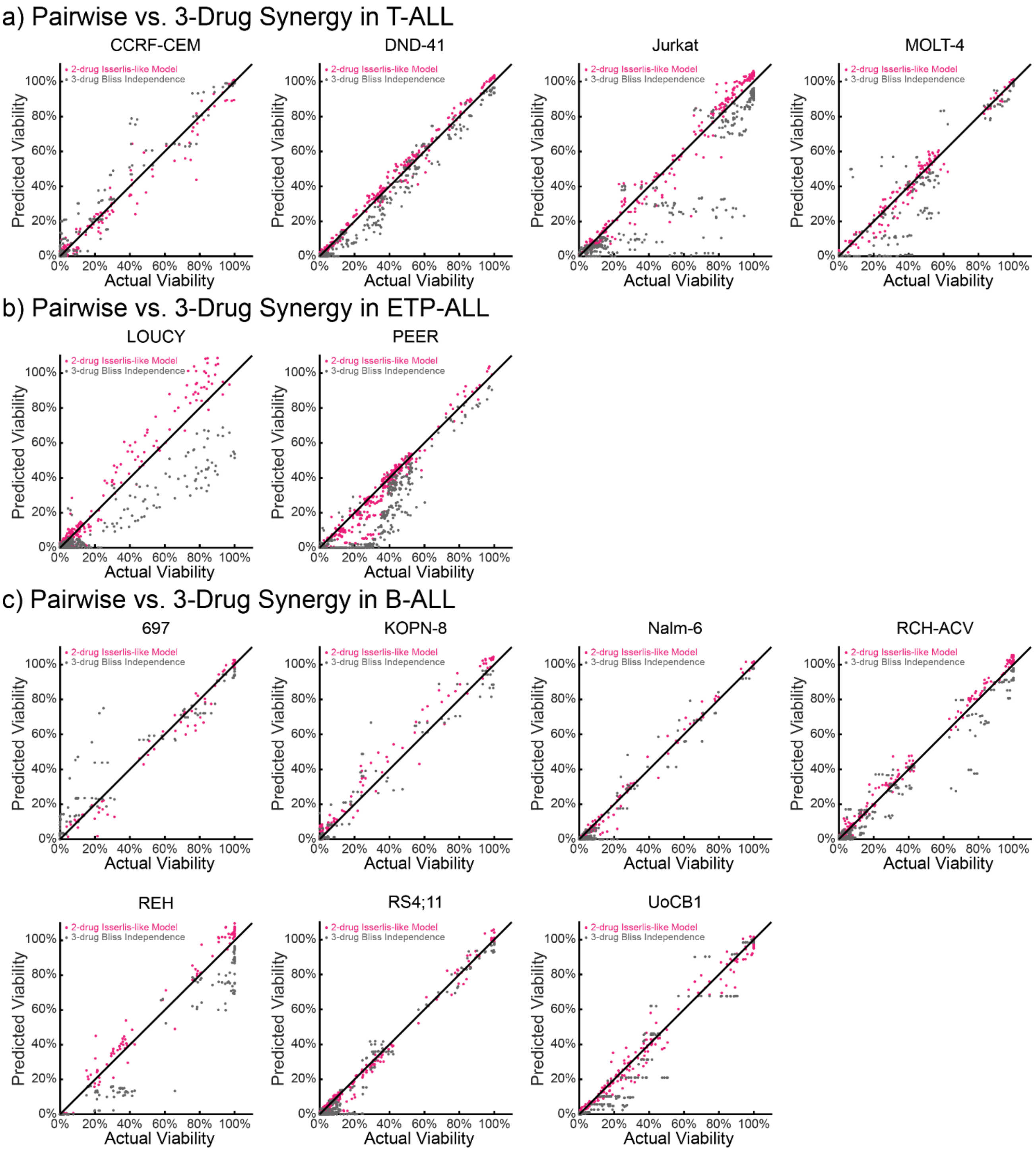


**SI Fig. 3. Drug synergy is predominantly attributable to pairwise rather than triplet drug interactions.** The Isserlis-like model incorporating 2-drug interactions better predicts responses to 3-drug combinations in **(a)** T-ALL, **(b)** ETP-ALL and **(c)** B-ALL compared to the Bliss Independence model, indicating that resulting dose responses are primarily attributable to 2-drug interactions rather than emergent 3-drug synergy between MRX-2843, methotrexate and vincristine. Data in (a-c) represent experimentally observed viabilities plotted against model predictions among 3-drug combination responses observed in high-throughput screens.

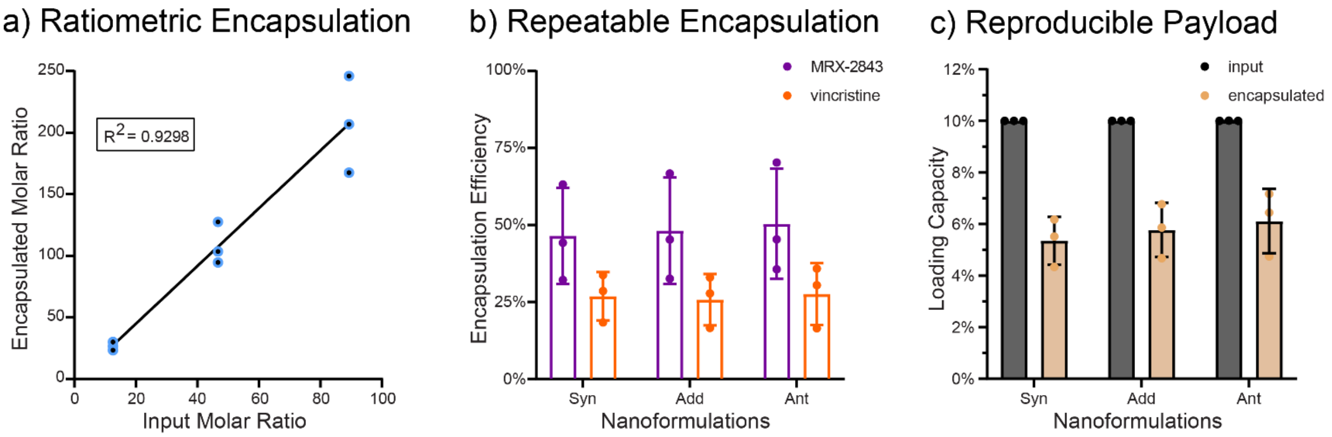


**SI Fig. 4. Co-encapsulation of MRX-2843 and vincristine is efficient and reproducible. a)** Calibration curve for ratiometric drug loading as measured by LC-MS. **b)** Encapsulation efficiencies, and **c)** loading capacities for Syn, Add, and Ant nanoparticles. The line in (a) represents best linear fit. Data in (a-c) represent individual measurements (dots) and mean values (bars) from 3 experiments.
